# Improved protein model quality assessment by integrating sequential and pairwise features using deep learning

**DOI:** 10.1101/2020.09.30.321661

**Authors:** Xiaoyang Jing, Jinbo Xu

## Abstract

**Motivation:** Accurately estimating protein model quality in the absence of experimental structure is not only important for model evaluation and selection, but also useful for model refinement. Progress has been steadily made by introducing new features and algorithms (especially deep neural networks), but accuracy of quality assessment (QA) is still not very satisfactory, especially local QA on hard protein targets.

**Results:** We propose a new single-model-based QA method ResNetQA for both local and global quality assessment. Our method predicts model quality by integrating sequential and pairwise features using a deep neural network composed of both 1D and 2D convolutional residual neural networks (ResNet). The 2D ResNet module extracts useful information from pairwise features such as model-derived distance maps, co-evolution information and predicted distance potential. The 1D ResNet is used to predict local (global) model quality from sequential features and pooled pairwise information generated by 2D ResNet. Tested on the CASP12 and CASP13 datasets, our experimental results show that our method greatly outperforms existing state-of-the-art methods. Our ablation studies indicate that the 2D ResNet module and pairwise features play an important role in improving model quality assessment.

**Availability and Implementation:** https://github.com/AndersJing/ResNetQA

## 1 Introduction

Significant progress has been made in computational protein structure prediction, especially template-free modeling (Wang *et al*., 2017; Xu, 2019; Senior *et al*., 2020; Kryshtafovych *et al*., 2019). To facilitate application of predicted 3D models, it is desirable to have an estimation of their local and global quality in the absence of experimental structures. Model quality assessment (QA) has also been used to assist protein model refinement (Heo *et al*., 2019; Park *et al*., 2019; Adiyaman and McGuffin, 2019; Hiranuma *et al*., 2020). Since CASP7 in 2006 (Cozzetto *et al*., 2007), many model QA methods have been developed, but the accuracy of local QA is still not very satisfactory, especially when all models under consideration are generated by an individual tool.

There are two major types of protein model QA methods: consensus (or clustering) methods and single-model methods (Won *et al*., 2019). Some methods are the combination of these two. Consensus methods mainly depend on clustering or comparison of models of a protein target. They work well when one protein has many models generated by different predictors, e.g., in CASPs. In the case there are very few models built by a single tool for a protein (which is often true in real-world application), single-model QA methods are needed, which predicts quality of a model using only its own information (Olechnovič and Venclovas, 2017; Uziela *et al*., 2017; Hurtado *et al*., 2018; Derevyanko *et al*., 2018; Karasikov *et al*., 2019). Most single-model QA methods extract some features from a decoy model and then map the features to a quality score using statistical or machine learning methods. A variety of features have been studied such as physical features, statistical features, local structural features (secondary structure and solvent accessibility), and sequence features (amino acid sequence and sequence profile). There are also a few quasi single-model methods that estimate model quality by comparing it to a small number of reference models generated by a small set of popular tools (Jing *et al*., 2016; Maghrabi and McGuffin, 2017). The quality of a protein model can be measured at residue/atom level (i.e., local quality) and at model level (i.e., global quality), which are referred as local and global quality assessment, respectively (Won *et al*., 2019; Cheng *et al*., 2019). Local quality is valuable for local structure error evaluation and model refinement, while global quality is valuable for model ranking and selection.

This paper focuses on single-model-based local and global QA. Although single-model QA is challenging, recently some progress has been made by using new features and deep learning. For example, ProQ3D uses a multi-layer perceptron to predict model quality from carefully-curated features (Uziela *et al*., 2017). ProQ4 predicts global QA by employing 1D fully convolutional neural network (CNN) and transfer learning (Hurtado *et al*., 2018). CNNQA applies 1D CNN to predict local and global model quality from sequential features, Rosetta energy terms, and knowledge-based potentials (Hou *et al*., 2019). QDeep (Shuvo *et al*., 2020) uses an ensemble of stacked 1D CNNs to predict quality from predicted distance information and some similar sequential features used by CNNQA. These methods mainly use coarse-grained structure representation and thus, may not perform well on local QA. 3D convolutional neural networks (Derevyanko *et al*., 2018; Pagès *et al*., 2019) or graph convolutional neural networks (Baldassarre *et al*., 2019; Sanyal *et al*., 2020; Igashov *et al*., 2020) are used to learn fine-grained structure representation, but these QA methods may not generalize well to protein models not extensively refined. One important issue with all these methods is that they do not take full advantage of inter-residue or inter-atom distance information which has greatly improved protein structure prediction recently (Zhu *et al*., 2018; Xu, 2019; Greener *et al*., 2019; Senior *et al*., 2020).

We propose a new single-model QA method ResNetQA (a ResNet-based QA method) that may greatly improve protein model QA, by using a deep 2D dilated residual network (ResNet) to explicitly extract useful information from pairwise features such as model-based distance matrices, predicted distance potentials and co-evolution information. These pairwise features may encode main structural information without introducing noise from side-chain or hydrogen atoms. Our method also uses a 1D deep ResNet to map sequential features (and pooled pairwise information derived from 2D ResNet) to local and global model quality. Further, to reduce bias introduced by a small training dataset, we train our deep model using a large set of decoy models of more than 14,000 proteins, in addition to CASP and CAMEO models. In particular, we built both template-based and template-free 3D models for ~14,000 proteins randomly selected from the CATH dataset (Dawson *et al*., 2017) using our in-house structure prediction software RaptorX. Our experimental results show that our method outperforms recently-developed single-model methods on both local and global QA.

## 2 Materials and methods

### 2.1 Overview

Figure 1 shows the overall architecture of our deep network, which mainly consists of one 2D ResNet module and one 1D ResNet module. The 2D ResNet module extracts information from pairwise features of shape L*L*N_2_ (where L is the sequence length and N_2_ is the dimension of pairwise features), which are derived from multiple sequence alignment (MSA) of a protein and its 3D models. This module outputs a high-level 2D feature map of shape L*L*C (C is the channel size), which is then converted to two 1D feature maps of shape L*C by mean pooling along row and column, respectively, and fed into the 1D ResNet module together with the original sequential features. The output of the 1D ResNet module is used to predict local and global model quality. To predict local quality, one fully-connected layer and one sigmoid layer are employed at each residue. To predict global quality, the output of the 1D ResNet module is converted to one vector of length (2C+N_1_) by mean pooling and fed into one fully-connected layer and one sigmoid layer.

**Fig. 1.**
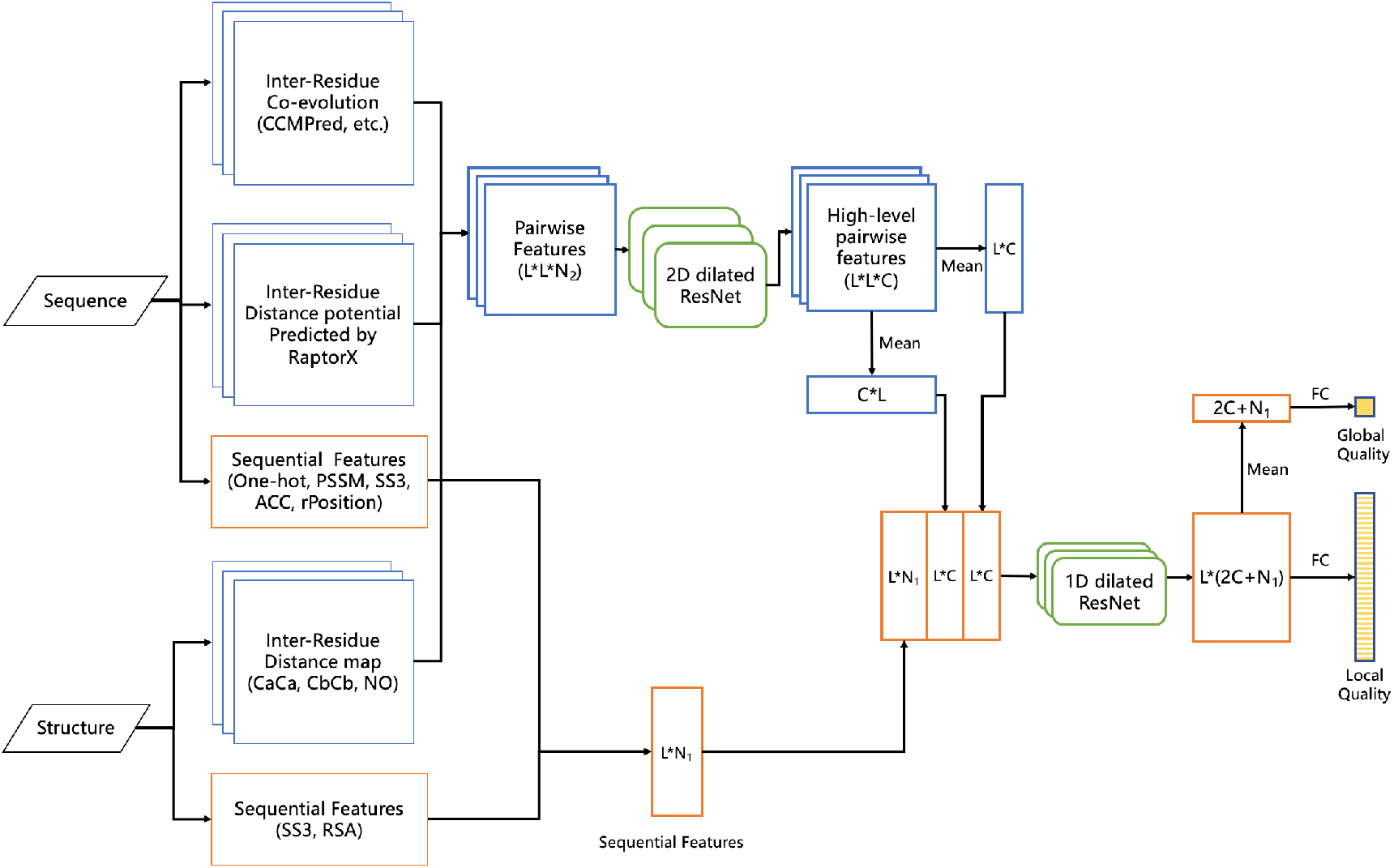
The overall architecture of our deep network for protein model local and global quality assessment. Meanwhile, L is the sequence length, C the channel size of the final 2D ResNet layer, N_1_ the dimension of sequential features, and N_2_ the dimension of pairwise features.

### 2.2 Datasets

We train and test our method using protein models from three sources: CASP, CAMEO and CATH. The CASP models were downloaded from http://predictioncenter.org/download_area/. The CAMEO (Haas *et al*., 2013) models are downloaded from https://www.cameo3d.org/sp/ and released between January 13, 2018 and September 14, 2019. Approximately 14,000 CATH domains (sequence identity <40%) are randomly selected from the CATH database (Dawson *et al*., 2017), and on average 15 template-based and template-free models are built for each domain using our in-house software RaptorX. These models are split into three datasets (training, validation and test) so that all models of one specific protein belong to only one dataset. The detailed information of our data is shown in the Supplementary Table S1.

The 3D models in CASP7-11 and CAMEO and the 3D models built for the 14,000 CATH domains are used as training and validation data. The 3D models in CASP12 and CASP13 are used as the test data. We remove their 3D models of proteins in the training and validation sets that share more than 25% sequence identity or have an BLAST E-value ≤0.001 (Altschul *et al*., 1997) with any CASP12 and CASP13 targets. As a result, there are 16450 proteins in our training and validation sets, among which ~5% are randomly selected to form the validation set. As a result, there are 15655 proteins with 364343 decoys in the training set and 795 proteins with 20483 decoys in the validation set. Since some proteins may have many more models than the others, at each training epoch we sample 320,000 decoys to train our deep model so that each protein has a similar number of models.

Only those CASP12 and CASP13 targets with publicly available experimental structures are used to test our method. As a result, there are 64 CASP12 targets with 9423 decoy models and 76 CASP13 targets with 11371 models in our test set. The 3D models in the CASP QA category are released in two stages. Since the decoy models in stage 1 are only used to check whether a method is a single-model method or a consensus method by comparing their predictions with those of stage 2 (Won *et al*., 2019), we mainly evaluate our method using the decoy models released at stage 2.

### 2.3 Feature extraction

From each protein sequence, we run HHblits (Remmert *et al*., 2012) to build its multiple sequence alignment (MSA) and then derive three types of features: sequential features, coevolution information and predicted distance potentials. Sequential features include: one-hot encoding of primary sequence (i.e., each residue is encoded as a binary vector of 21 entries indicating its amino acid type), rPosition (the relative position of a residue in a sequence calculated as *i/L* where *i* is the residue index), PSSM (position-specific scoring matrix derived from MSA), SS3 (3-state secondary structure predicted by RaptorX-Property (Wang *et al*., 2016)), and ACC (solvent accessibility predicted by RaptorX-Property (Wang *et al*., 2016)). Coevolution information includes the output generated by CCMPred (Seemayer *et al*., 2014) and raw and APC-corrected mutual information (MI). Distance potentials is derived from distance distribution predicted by RaptorX-Contact (Xu, 2019) from MSA. Only C_β_-C_β_ distance potential is used, and distance is discretized into 14 bins: <4, 4-5, 5-6, …, 14-15, 15-16, >16. From each protein model, we derive the following structural features: 1) secondary structure (SS3) and relative solvent accessibility (RSA) calculated by DSSP (Kabsch and Sander, 1983) from a 3D model; and 2) distance maps of three atom pairs (C_α_C_α_, C_β_C_β_ and NO) calculated from a 3D model. Supplementary Table S2 summarizes all these features.

### 2.4 Deep neural network architecture and training

As shown in Figure 1, our deep model mainly consists of 1D and 2D dilated residual neural networks (ResNet) (He *et al*., 2016). A gate block is used to connect pairwise input features to the 2D ResNet. It is composed of one 2D convolutional layer, one instance norm layer (Ulyanov *et al*., 2017) and one ELU (exponential linear unit) activation layer. As shown in Figure 2, one ResNet block consists of 2 instance norm layer, 2 convolutional layer, 2 ELU activation layer and 1 dropout layer. In a ResNet block there is a shortcut connecting its input to the output of the second convolutional layer. In order to capture a broader context, a dilation ratio of 2 is used in a 2D convolutional layer. In summary, our deep model contains one gate block, 10 2D ResNet blocks with 64 filters of size 5*5, 8 1D ResNet blocks with 180 filters of size 5. That is, in total there are 21 2D convolutional layers and 16 1D convolutional layers in our deep model. We have also tested more ResNet blocks, but not observed significant improvement.

**Fig. 2.**
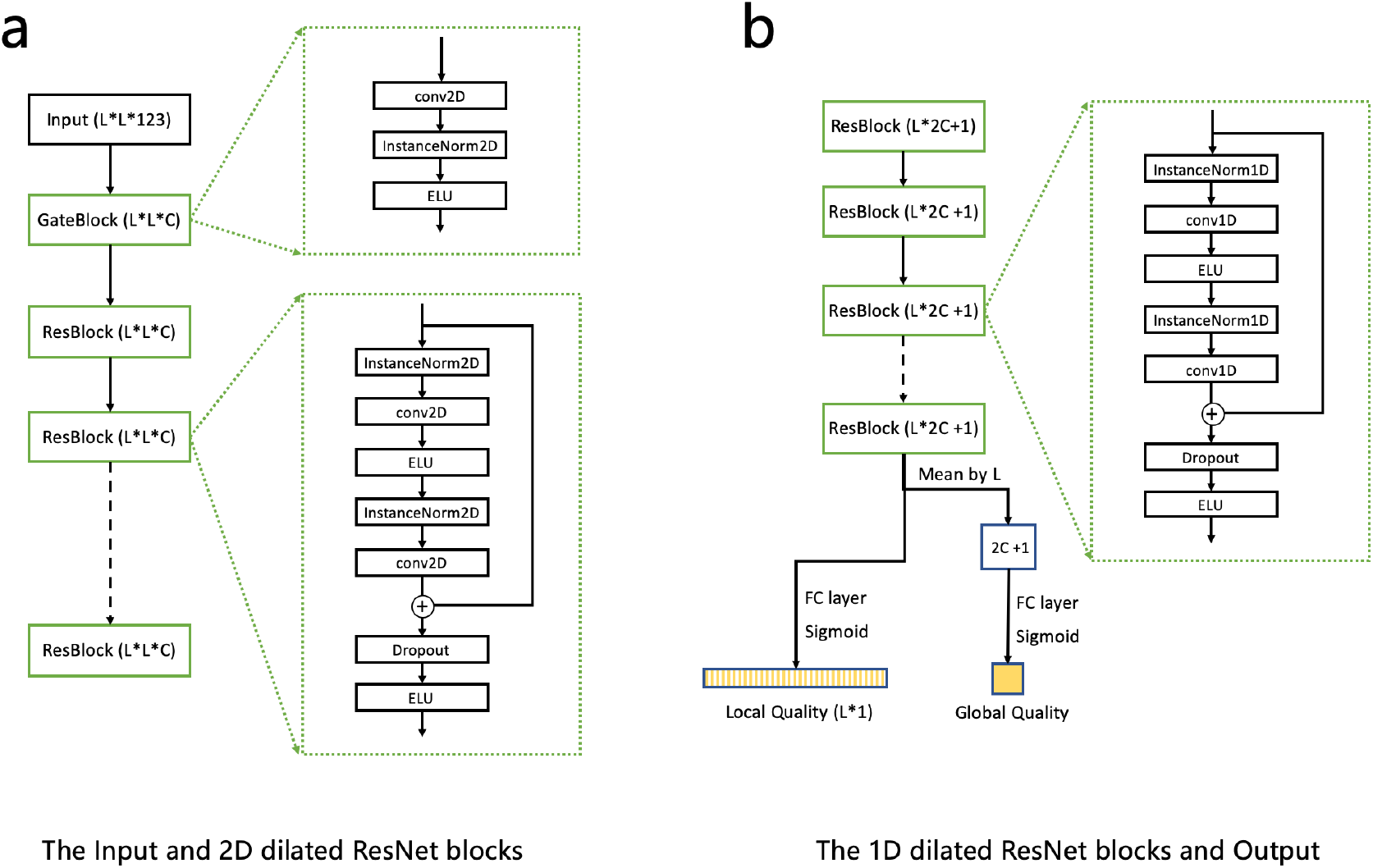
The architecture of a gate block, a 2D dilated ResNet block and a 1D dilated ResNet block.

A protein may not fit into the limited memory of a graphics processing unit (GPU). To deal with this, a sequence segment of length 350 (and its corresponding sequential and pairwise features) is randomly sampled when a protein has more than 350 residues. We implement our method with PyTorch (Paszke *et al*., 2019) and train it using the Adam optimizer with parameters: β1=0.9 and β2=0.999. We set the initial learning rate to 0.0001 and divide it by 2 every 3 epochs. Each minibatch has 16 3D models. Since the training dataset is very large, the training procedure usually converges within 10 epochs. The deep model with minimum loss on the validation data is selected as our final deep model.

For local QA, our deep model predicts a residue-wise S-score defined by 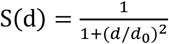 where d is the distance deviation of one C_α_ atom from its position in the experimental structure calculated by LGA (Zemla, 2003). Here we set d_0_ to 3.8Å instead of 5.0Å to yield accurate prediction for small d. We convert predicted S-score to predicted distance error (or deviation) by the inverse function of S(d). For global QA, our deep model predicts GDT_TS (Global Distance Test Total Score). The loss of our deep model is the MSE (Mean Square Error) between predicted local (or global) quality and its ground truth. Our deep network is trained to simultaneously predict local and global quality and equal weight is used to combine their losses.

### 2.5 Evaluation Metrics

We employ several widely-used metrics (Won *et al*., 2019) to evaluate the performance of our method. To evaluate local QA, we use PCC, ASE and AUC.

- PCC is the Pearson correlation coefficient between predicted local quality score S-score and its ground truth. All models of a specific protein are pooled together when calculating PCC of local QA.
- ASE: ASE is the average residue-wise S-score similarity defined as 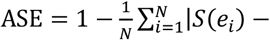 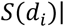, where N is the number of residues, S(*e*_*i*_) is the S-score derived from predicted distance deviation and S(*d*_*i*_) is the S-score derived from the true distance deviation produced by LGA (Zemla, 2003). Following CASP, here the S-score is defined by 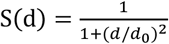 where d_0_ = 5.0Å.
- AUC: The AUC (Area Under Curve) assesses how well the predicted score may distinguish accurate residues from inaccurate ones, where an accurate residue is the one with Cα atom deviates from its experimental position by no more than 3.8Å. AUC is the area under the ROC curve, which plots the TP (true positive rate) against the FP (false positive rate) in the prediction of accurate/inaccurate residues by varying the cut-off score for distinguishing accurate/inaccurate residues. For global QA, we use PCC, Diff and Loss as metrics. Global metrics are calculated and averaged at the protein level.
- PCC: The Pearson correlation coefficient between predicted global quality scores and ground truth.
- Diff: the mean absolute difference between predicted global quality and ground truth.
- Loss: The absolute quality difference (measured by GDT-TS) between the best model predicted by a QA method and the real best model.

## 3 Results and discussion

### 3.1 Performance on CASP12 and CASP13 datasets

We compare our method with some single-model methods ranked well in CASP12 and CASP13 (Won *et al*., 2019; Kryshtafovych *et al*., 2018), e.g., ProQ3, Wang4, ProQ2, ZHOU-SPARKS-X and VoroMQA in CASP12 and ProQ4, VoroMQA-A, VoroMQA-B, ProQ3D and ProQ3 in CASP13. Their local and global quality estimations are downloaded from the CASP official website.

Table 1 lists the performances on the CASP12 and CASP13 stage 2 datasets. As shown in the table, our method significantly outperforms others on both datasets in terms of most evaluation metrics. On the CASP12 dataset, ProQ3 is slightly better than the other four methods on both local and global QA, while our method greatly outperforms ProQ3. When local QA is evaluated, our method is ~29% better than ProQ3 in terms of PCC (0.5866 vs. 0.4542), ~15% better in terms of ASE (0.8515 vs. 0.7409) and ~7% better in terms of AUC (0.8058 vs. 0.7517). When global QA is evaluated, our method is >20% better than ProQ3 in terms of both PCC (0.8109 vs. 0.6552) and Diff (0.0785 vs. 0.1104). On the CASP13 dataset, our method has similar advantage over the others. When local QA is evaluated, our method is ~30% better than ProQ3D in terms of PCC (0.5539 vs. 0.4230) and ~15% better in terms of ASE (0.8373 vs. 0.7312). When global QA is evaluated, our method is ~25% better than ProQ3D in terms of PCC (0.8157 vs. 0.6532) and ~19% better in terms of Diff (0.0861 vs. 0.1060). In terms of Loss, our method is not much better than ProQ3 and ProQ3D when the mean value of Loss is compared. However, our method has a smaller standard deviation of Loss, which indicates that our method is more robust across all tested targets.

**Table 1.** Performances of single-model methods on local and global QA

| Dataset | Method<sup>1</sup> | Local |  |  | Global |  |  |
| --- | --- | --- | --- | --- | --- | --- | --- |
| | | PCC↑ | ASE↑ | AUC↑ | PCC↑ | Diff↓ | Loss↓ |
| CASP12<br>Stage 2 | **ResNetQA** | **0.5866** | **0.8515** | **0.8058** | **0.8109** | **0.0785** | **0.0612**($\pm 0.0665^2$) |
| | ProQ3 | 0.4542 | 0.7409 | 0.7517 | 0.6552 | 0.1104 | 0.0615($\pm 0.0649$) |
| | ProQ2 | 0.4364 | 0.6928 | 0.7434 | 0.6108 | 0.1338 | 0.0707($\pm 0.0682$) |
| | Wang4 | 0.4128 | 0.7459 | 0.7229 | 0.6206 | 0.1282 | 0.1162($\pm 0.1327$) |
| | VoroMQA | 0.4099 | 0.7122 | 0.7254 | 0.6071 | 0.1668 | 0.0844($\pm 0.0918$) |
| | ZHOU-SPARKS-X | 0.3873 | 0.7655 | 0.7100 | 0.6757 | 0.1369 | 0.0779($\pm 0.0756$) |
| CASP13<br>Stage 2 | **ResNetQA** | **0.5539** | **0.8373** | **0.7901** | **0.8157** | **0.0861** | **0.0844**($\pm 0.0713$) |
| | ProQ3D | 0.4230 | 0.7312 | 0.7384 | 0.6532 | 0.1060 | 0.0852($\pm 0.0891$) |
| | ProQ3 | 0.4134 | 0.7205 | 0.7455 | 0.5921 | 0.1198 | 0.0898($\pm 0.0918$) |
| | ProQ4 | 0.3873 | 0.6100 | 0.7235 | 0.7190 | 0.1371 | 0.0871($\pm 0.0908$) |
| | VoroMQA-A | 0.3872 | 0.6704 | 0.7238 | 0.6439 | 0.1548 | 0.1095($\pm 0.1442$) |
| | VoroMQA-B | 0.3831 | 0.6705 | 0.7214 | 0.6189 | 0.1568 | 0.0995($\pm 0.1315$) |

1. Methods sorted in decreasing order by PCC of local QA.
2. The standard deviation on the Loss metric.

### 3.2 Performance on CASP12 and CASP13 FM targets

The structure of a template-free modeling (FM) target may be harder to predict and the predicted 3D models may have very different quality (Kinch *et al*., 2019), which makes QA harder. Here we evaluate our method using the 3D models of CASP12 and CASP13 FM targets. One target is FM if it contains at least one FM evaluation unit by the CASP official definition (Abriata *et al*., 2018; Kinch *et al*., 2019).

As shown in Table 2, all methods have worse performance than what is shown in Table 1. However, our method shows a larger advantage over the other methods. When local QA on the CASP12 FM targets is evaluated, our method is better than the 2^nd^ best method ProQ3 by ~57% in terms of PCC (0.4445 vs. 0.2830), by ~28% in terms of ASE (0.8930 vs. 0.6951) and by ~12% in terms of AUC (0.7487 vs. 0.6696). When global QA is evaluated, our method is ~34% better than the 2^nd^ best method in terms of PCC (0.7213 vs. 0.5381) and ~40% better in terms of Diff (0.0584 vs. 0.0978). On the CASP13 FM targets, compared with the ProQ3D, the advantage of our method on local QA is: ~60% on PCC, ~25% on ASE, and ~8% on AUC. On global QA, our method is better by ~42% on PCC and by ~12% on Diff, respectively. The Loss metric on global QA varies a lot. ProQ4 performs slightly better than our method in terms of the mean value of Loss, but our method has a much smaller standard deviation.

**Table 2.**
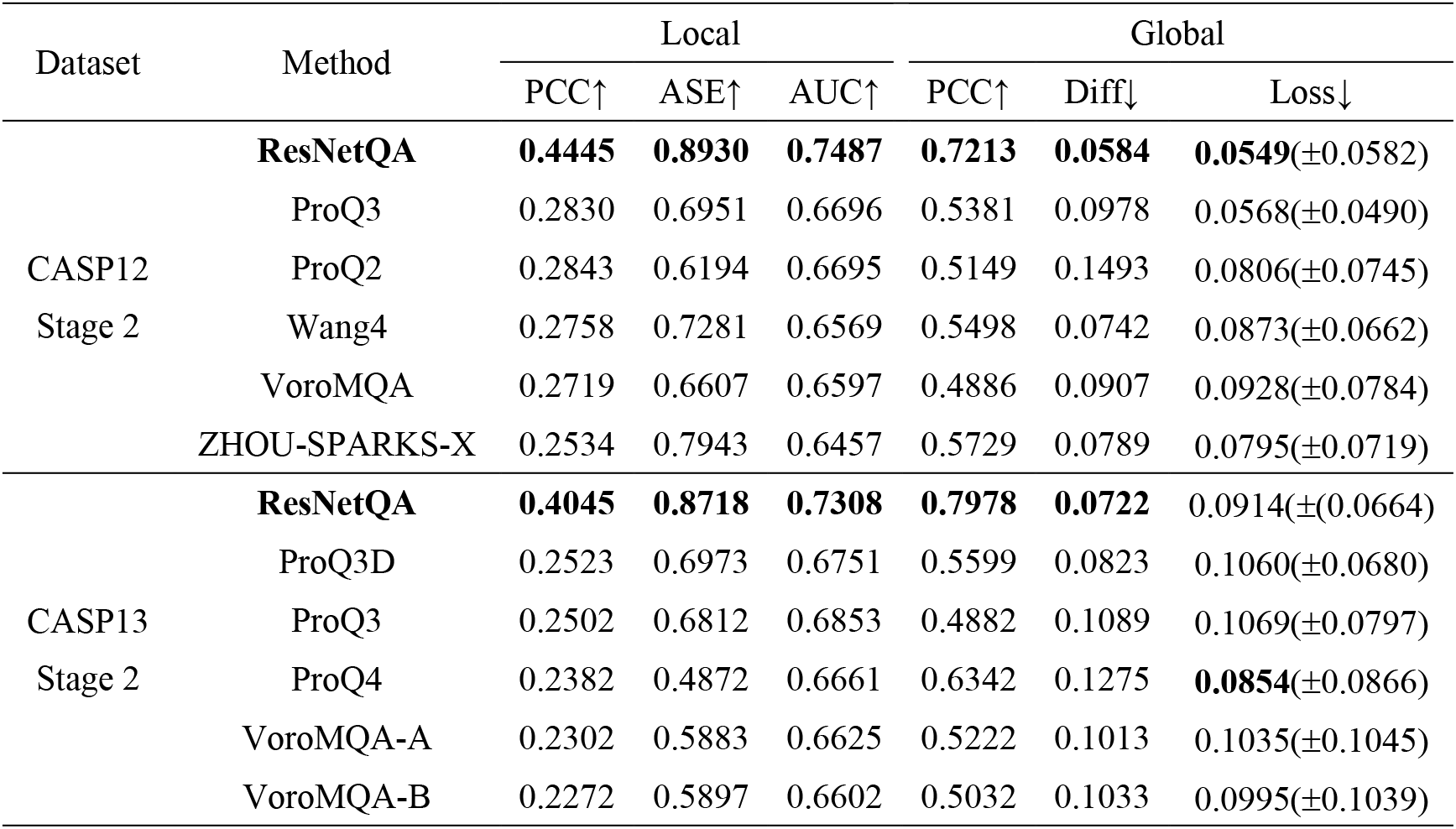
Performances of single-model methods on CASP12 and CASP13 FM targets

### 3.3 Comparison with other deep learning methods

We compare our method with three leading single-model QA methods: ProQ4 (Hurtado *et al*., 2018), CNNQA (Hou *et al*., 2019) and QDeep (Shuvo *et al*., 2020). Meanwhile, ProQ4 performed very well in CASP13 and QDeep predicts only global quality. All these methods are built upon 1D CNN and thus, cannot make a very good use of pairwise features. Both ProQ4 and CNNQA use only sequential features. Although QDeep indeed uses predicted inter-residue distance information, its simple way of converting predicted distance information to sequential features is not as effective as the 2D ResNet used by our method. CNNQA and QDeep use energy potentials as features, but our method does not. There may be some valuable information in energy potential and we will try out this in future work.

We evaluate our method on the same set of 40 CASP12 targets as QDeep and CNNQA were evaluated. We run ProQ4 locally with default parameters and convert its prediction to distance error by the S-function with d_0_=5.0. As shown in Table 3, our method outperforms these three methods in terms of 5 metrics.

**Table 3.** Performances of deep learning QA methods on 40 CASP12 targets^1^

| Method | Local |  |  | Global |  |  |
| --- | --- | --- | --- | --- | --- | --- |
| | PCC↑ | ASE↑ | AUC↑ | PCC↑ | Diff↓ | Loss↓ |
| **ResNetQA** | **0.5730** | **0.8526** | **0.7995** | **0.7990** | **0.0746** | 0.0639($\pm 0.0723$) |
| ProQ4 | 0.3884 | 0.7581 | 0.7085 | 0.6617 | 0.1419 | 0.0670($\pm 0.0691$) |
| CNNQA | -- | 0.7814 | -- | 0.6270 | -- | 0.0854(--) |
| QDeep | -- | -- | -- | 0.7400 | 0.1053 | **0.0510($\pm 0.0480$)** |

1. The 40 CASP12 targets include target T0865 which was canceled later by CASP12. Here we still use it to be consistent with CNNQA and QDeep.

### 3.4 Contribution of different input features

We have trained three extra models by excluding 1) mutual information and coevolution information produced by CCMPred; 2) predicted and model-derived secondary structures and solvent accessibilities; or 3) predicted distance potentials. As a control, we also trained a 1D deep model of 16 1D ResNet blocks without using any 2D ResNet blocks to predict model quality mainly from sequential features. To feed predicted distance potential to this 1D deep model, we apply mean pooling to distance potential at each residue. That is, we calculate the average of all pairwise potentials involving a specific residue and use the average as an extra feature of this residue. By this way, our 1D deep model can also make use of pairwise information, although not very effectively.

Figure 3 shows the PCC of these deep models on local QA. Their detailed performances are available in Supplementary Table S3. It is not surprising that the 2D deep model built with all features performs the best. The 1D deep model performs the worst, which implies that the 2D ResNet module is very important. Compared to the deep model built with all features, the 2D deep model using co-evolution information but not predicted distance potential is about 9% worse on the CASP12 dataset and ~11% worse on the CASP13 dataset. The 2D deep model without predicted and model-derived secondary structure and solvent accessibilities is about ~4% worse on the CASP12 dataset and ~2% worse on the CASP13 dataset. The 2D deep model using predicted distance but not co-evolution information is slightly worse than the deep model built with all features, because predicted distance potential already encodes most co-evolution information.

**Fig. 3.**
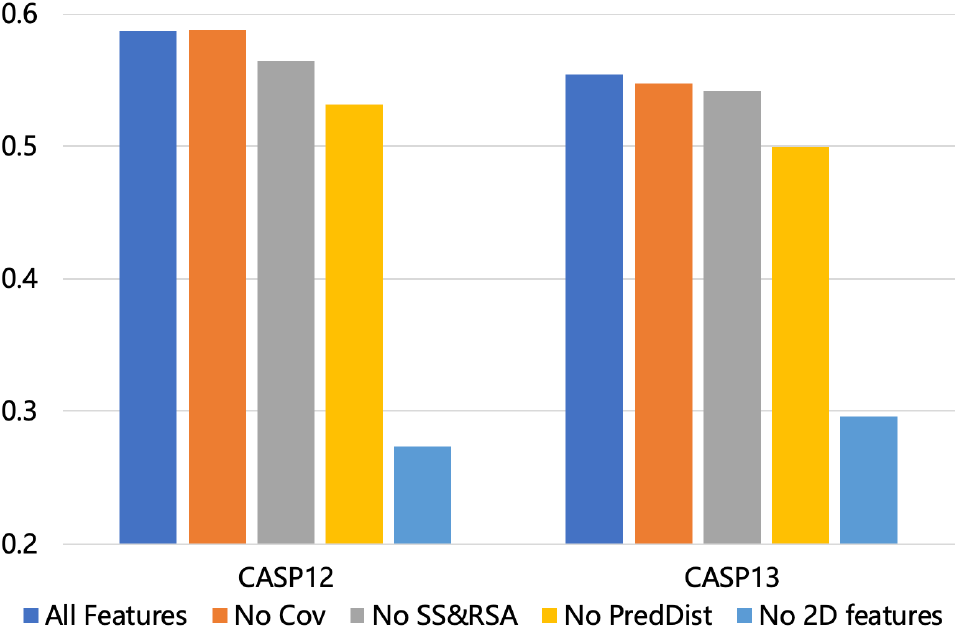
Performance (measured by PCC on local QA) of deep models built with different features.

### 3.5 Contribution of the CATH dataset

Many QA methods are trained by the 3D models of only hundreds of proteins in CASP and CAMEO, which have limited coverage of the whole protein universe (Bateman *et al*., 2017). In contrast, protein structural property prediction models (Wang *et al*., 2016) and contact/distance prediction models (Xu, 2019; Senior *et al*., 2020) usually are trained by thousands and even tens of thousands proteins. To reduce bias incurred by a small number of training proteins and improve generalization capability, we built both template-based and template-free models for about 14,000 proteins randomly selected from the CATH dataset (Dawson *et al*., 2017) using our in-house structure prediction software RaptorX and use these models as training data. Here we compare the performance of two deep models with exactly the same architecture and the same input features, but trained by different data. One is trained using all models built on the CASP, CAMEO and CATH datasets and the other is trained using only the CASP and CAMEO models.

Figure 4 shows a head-to-head comparison of these two deep models in terms of PCC on local QA. Their detailed performance is available in Supplementary Table S4. It is clear that the deep model trained by all 3D models works much better. On the CASP12 dataset, the deep model trained using all 3D models outperforms that trained without the CATH data by about 7.6% (0.5866 vs. 0.5450). On the CASP13 data, the advantage is about 5.0% (0.5539 vs. 0.5276). The result suggests that the decoy models built for CATH data is valuable for improving protein model quality assessment.

**Fig. 4.**
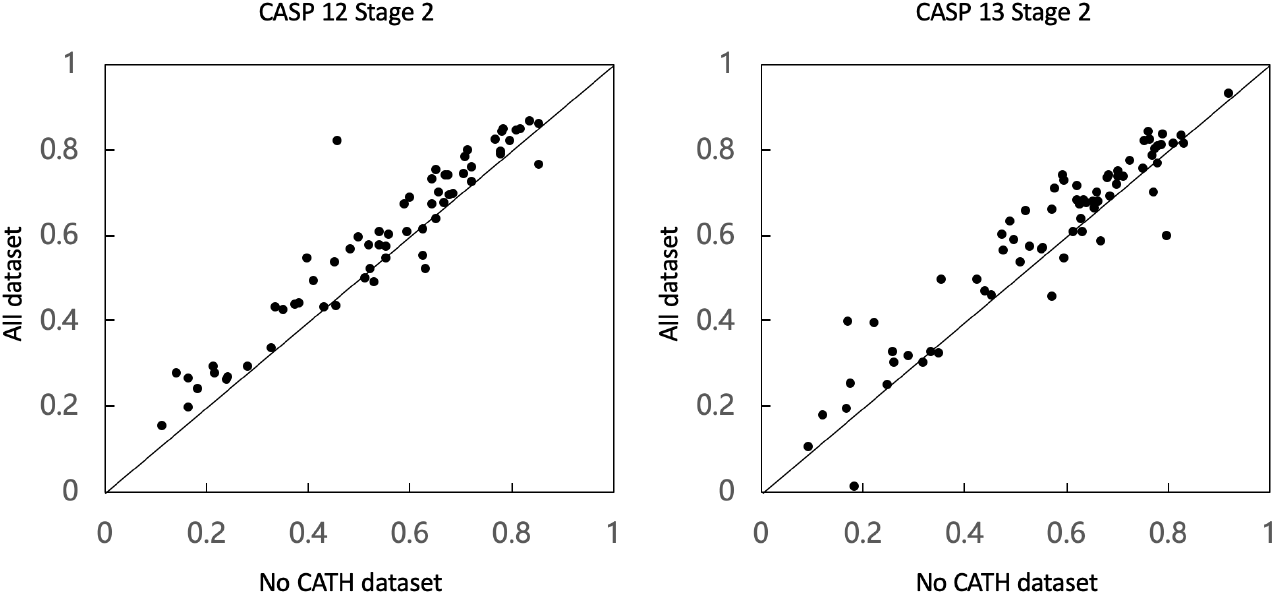
Head-to-head performance (measured by PCC on local QA) comparison of the two deep models trained with and without the CATH data.

## 4 Conclusion

In this paper we have presented a new single-model QA method ResNetQA for both local and global protein model quality estimation. Our method predicts model quality by integrating a variety of sequential and pairwise features using a deep network composed of both 1D and 2D ResNet blocks. The 2D ResNet blocks extracts useful information from pairwise features and the 1D ResNet blocks predict quality from sequential features and pairwise information produced by the 2D ResNet module. Our method differs from existing ones mainly by the 2D ResNet module and a larger training set. Our test results on the CASP12 and CASP13 datasets show that our method significantly outperforms existing state-of-the-art methods, especially on hard targets. Our ablation studies confirm that the 2D ResNet module and pairwise features are very important for the superior performance of our method. Recent research shows that inter-residue orientation is valuable for protein structure prediction (Yang *et al*., 2020). QA may be further improved by incorporating this kind of information.

## References

Abriata,L.A. et al. (2018) Definition and classification of evaluation units for tertiary structure prediction in CASP12 facilitated through semi-automated metrics. Proteins: Structure, Function, and Bioinformatics, 86, 16–26.

Adiyaman,R. and McGuffin,L.J. (2019) Methods for the Refinement of Protein Structure 3D Models. International Journal of Molecular Sciences, 20, 2301.

Altschul,S.F. et al. (1997) Gapped BLAST and PSI-BLAST: a new generation of protein database search programs. Nucleic acids research, 25, 3389–3402.

Baldassarre,F. et al. (2019) GraphQA: Protein Model Quality Assessment using Graph Convolutional Network.

Bateman,A. et al. (2017) UniProt: the universal protein knowledgebase. Nucleic Acids Res, 45, D158–D169.

Cheng,J. et al. (2019) Estimation of model accuracy in CASP13. Proteins: Structure, Function, and Bioinformatics, 87, 1361–1377.

Cozzetto,D. et al. (2007) Assessment of predictions in the model quality assessment category. Proteins: Structure, Function, and Bioinformatics, 69, 175–183.

Dawson,N.L. et al. (2017) CATH: an expanded resource to predict protein function through structure and sequence. Nucleic Acids Res, 45, D289–D295.

Derevyanko,G. et al. (2018) Deep convolutional networks for quality assessment of protein folds. Bioinformatics, 34, 4046–4053.

Greener,J.G. et al. (2019) Deep learning extends de novo protein modelling coverage of genomes using iteratively predicted structural constraints. Nature Communications, 10, 1–13.

Haas,J. et al. (2013) The Protein Model Portal--a comprehensive resource for protein structure and model information. Database (Oxford), 2013, bat031.

He,K. et al. (2016) Deep Residual Learning for Image Recognition., pp. 770–778.

Heo,L. et al. (2019) Driven to near-experimental accuracy by refinement via molecular dynamics simulations. Proteins: Structure, Function, and Bioinformatics, 87, 1263–1275.

Hiranuma,N. et al. (2020) Improved protein structure refinement guided by deep learning based accuracy estimation. bioRxiv, 2020.07.17.209643.

Hou,J. et al. (2019) Deep convolutional neural networks for predicting the quality of single protein structural models. bioRxiv, 590620.

Hurtado,D.M. et al. (2018) Deep transfer learning in the assessment of the quality of protein models. arXiv:1804.06281 [q-bio].

Igashov,I. et al. (2020) VoroCNN: Deep convolutional neural network built on 3D Voronoi tessellation of protein structures. bioRxiv, 2020.04.27.063586.

Jing,X. et al. (2016) Sorting protein decoys by machine-learning-to-rank. Scientific Reports, 6, 1–11.

Kabsch,W. and Sander,C. (1983) Dictionary of protein secondary structure: Pattern recognition of hydrogen-bonded and geometrical features. Biopolymers, 22, 2577–2637.

Karasikov,M. et al. (2019) Smooth orientation-dependent scoring function for coarse-grained protein quality assessment. Bioinformatics, 35, 2801–2808.

Kinch,L.N. et al. (2019) CASP13 target classification into tertiary structure prediction categories. Proteins: Structure, Function, and Bioinformatics, 87, 1021–1036.

Kryshtafovych,A. et al. (2018) Assessment of model accuracy estimations in CASP12. Proteins: Structure, Function, and Bioinformatics, 86, 345–360.

Kryshtafovych,A. et al. (2019) Critical assessment of methods of protein structure prediction (CASP)—Round XIII. Proteins: Structure, Function, and Bioinformatics, 87, 1011–1020.

Maghrabi,A.H.A. and McGuffin,L.J. (2017) ModFOLD6: an accurate web server for the global and local quality estimation of 3D protein models. Nucleic Acids Res, 45, W416–W421.

Olechnovič,K. and Venclovas,Č. (2017) VoroMQA: Assessment of protein structure quality using interatomic contact areas. Proteins: Structure, Function, and Bioinformatics, 85, 1131–1145.

Pagès,G. et al. (2019) Protein model quality assessment using 3D oriented convolutional neural networks. Bioinformatics, btz122.

Park,H. et al. (2019) High-accuracy refinement using Rosetta in CASP13. Proteins: Structure, Function, and Bioinformatics, 87, 1276–1282.

Paszke,A. et al. (2019) PyTorch: An Imperative Style, High-Performance Deep Learning Library. In, Wallach,H. et al. (eds), Advances in Neural Information Processing Systems 32. Curran Associates, Inc., pp. 8026–8037.

Remmert,M. et al. (2012) HHblits: lightning-fast iterative protein sequence searching by HMM-HMM alignment. Nature Methods, 9, 173–175.

Sanyal,S. et al. (2020) ProteinGCN: Protein model quality assessment using Graph Convolutional Networks. bioRxiv, 2020.04.06.028266.

Seemayer,S. et al. (2014) CCMpred--fast and precise prediction of protein residue-residue contacts from correlated mutations. Bioinformatics, 30, 3128–30.

Senior,A.W. et al. (2020) Improved protein structure prediction using potentials from deep learning. Nature, 577, 706–710.

Shuvo,M.H. et al. (2020) QDeep: distance-based protein model quality estimation by residue-level ensemble error classifications using stacked deep residual neural networks. bioRxiv, 2020.01.31.928622.

Ulyanov,D. et al. (2017) Instance Normalization: The Missing Ingredient for Fast Stylization. arXiv:1607.08022 [cs].

Uziela,K. et al. (2017) ProQ3D: improved model quality assessments using deep learning. Bioinformatics, 33, 1578–1580.

Wang,S. et al. (2017) Accurate De Novo Prediction of Protein Contact Map by Ultra-Deep Learning Model. PLOS Computational Biology, 13, e1005324.

Wang,S. et al. (2016) RaptorX-Property: a web server for protein structure property prediction. Nucleic Acids Res, 44, W430–W435.

Won,J. et al. (2019) Assessment of protein model structure accuracy estimation in CASP13: Challenges in the era of deep learning. Proteins: Structure, Function, and Bioinformatics, 87, 1351–1360.

Xu,J. (2019) Distance-based protein folding powered by deep learning. PNAS, 116, 16856–16865.

Yang,J. et al. (2020) Improved protein structure prediction using predicted interresidue orientations. PNAS, 117, 1496–1503.

Zemla,A. (2003) LGA: A method for finding 3D similarities in protein structures. Nucleic acids research, 31, 3370.

Zhu,J. et al. (2018) Protein threading using residue co-variation and deep learning. Bioinformatics, 34, i263–i273.

