## Supplementary information for "Improved protein model quality assessment by integrating sequential and pairwise features using deep learning"

Table S1. The number of protein targets and decoys of different datasets.

| Dataset | Data Source | #Targets | | #Decoys | |
|---|---|---|---|---|---|
| Training | CASP 7-11 | 435 | 15655 | 101153 | 364343 |
| | CAMEO | 1182 | | 49139 | |
| | CATH | 14038 | | 214051 | |
| Validation | CASP 7-11 | 27 | 795 | 6943 | 20483 |
| | CAMEO | 70 | | 2863 | |
| | CATH | 698 | | 10677 | |
| Test | CASP12 | 64 | | 9423 | |
| | CASP13 | 76 | | 11371 | |

Table S2. Summary of input features

| Type | Feature | Type | Shape |
|---|---|---|---|
| Derived from Sequence | One-hot encoding of sequence | Sequential | L*21 |
| | rPosition | Sequential | L*1 |
| | PSSM | Sequential | L*20 |
| | Predicted SS3 | Sequential | L*3 |
| | Predicted ACC | Sequential | L*3 |
| | co-evolution | Pairwise | L*L*4 |
| | Predicted distance potential | Pairwise | L*L*14 |
| Derived from model structure | SS3 | Sequential | L*3 |
| | RSA | Sequential | L*1 |
| | distance (CaCa, CbCb and NO) | Pairwise | L*L*3 |
| Total | | | Sequential: L*52<br>Pairwise: L*L*21 |

Table S3. Performances of deep models built with different features

| Dataset | Method | Local | | | Global | | |
|---|---|---|---|---|---|---|---|
| | | PCC↑ | ASE↑ | AUC↑ | PCC↑ | Diff↓ | Loss↓ |
| CASP12 Stage 2 | **ResNetQA** | 0.5866 | **0.8515** | 0.8058 | **0.8109** | 0.0785 | **0.0612** |
| | No Cov | **0.5874** | 0.8507 | **0.8063** | 0.8008 | **0.0758** | 0.0730 |
| | No SS&RSA | 0.5643 | 0.8302 | 0.7948 | 0.7788 | 0.0877 | 0.0733 |
| | No PredDist | 0.5319 | 0.8243 | 0.7847 | 0.7192 | 0.0957 | 0.0776 |
| | No 2D features | 0.2735 | 0.6965 | 0.7912 | 0.4046 | 0.1646 | 0.1192 |
| CASP13 Stage 2 | **ResNetQA** | **0.5539** | **0.8373** | **0.7901** | **0.8157** | **0.0861** | 0.0844 |
| | No Cov | 0.5474 | 0.8248 | 0.7889 | 0.8020 | 0.0897 | **0.0829** |
| | No SS&RSA | 0.5416 | 0.8057 | 0.7615 | 0.7771 | 0.0969 | 0.1068 |
| | No PredDist | 0.5000 | 0.7971 | 0.7501 | 0.7450 | 0.1057 | 0.0999 |
| | No 2D features | 0.2963 | 0.6981 | 0.6472 | 0.5119 | 0.1524 | 0.1636 |

Table S4. Performances of deep models trained with and without the CATH dataset

| Dataset | Method | Local | | | Global | | |
|---|---|---|---|---|---|---|---|
| | | PCC↑ | ASE↑ | AUC↑ | PCC↑ | Diff↓ | Loss↓ |
| CASP12 Stage 2 | **ResNetQA** | **0.5866** | **0.8515** | **0.8058** | **0.8109** | **0.0785** | **0.0612** |
| | No CATH | 0.5450 | 0.8425 | 0.7689 | 0.7520 | 0.0893 | 0.0996 |
| CASP13 Stage 2 | **ResNetQA** | **0.5539** | **0.8373** | **0.7901** | **0.8157** | **0.0861** | **0.0844** |
| | No CATH | 0.5276 | 0.8134 | 0.7487 | 0.7873 | 0.1211 | 0.0947 |
